# Deep kinematic inference affords efficient and scalable control of bodily movements

**DOI:** 10.1101/2023.05.04.539409

**Authors:** Matteo Priorelli, Giovanni Pezzulo, Ivilin Peev Stoianov

## Abstract

Performing goal-directed movements requires mapping goals from extrinsic (workspace-relative) to intrinsic (body-relative) coordinates and then to motor signals. Mainstream approaches based on Optimal Control realize the mappings by minimizing cost functions, which is computationally demanding. Instead, Active Inference uses generative models to produce sensory predictions, which allows a cheaper inversion to the motor signals. However, devising generative models to control complex kinematic chains like the human body is challenging. We introduce a novel Active Inference architecture that affords a simple but effective mapping from extrinsic to intrinsic coordinates via inference and easily scales up to drive complex kinematic chains. Rich goals can be specified in both intrinsic and extrinsic coordinates using attractive or repulsive forces. The proposed model reproduces sophisticated bodily movements and paves the way for computationally efficient and biologically plausible control of actuated systems.

## 1 Introduction

How do the brains of living systems support goal-directed movements implementing purposeful behavior? A standard assumption is that performing goal-directed actions requires two mappings reflecting perceptual and plant control processes [1]. First, it is necessary to map goals and desired movement trajectories specified in *extrinsic* coordinates (e.g., in Cartesian space) into movements specified in *intrinsic* coordinates (e.g., as joint angles), via a process called *inverse kinematics*. Second, it is necessary to map the intrinsic state into the forces needed to move all the muscles of the body, via a process called *inverse dynamics*.

These two inversions (kinematic and dynamic) are computationally challenging. While computing the extrinsic representation of a given joint configuration is easy as the geometric mapping – the so-called *direct kinematics* – is univocal, finding the joint angles that correspond to a particular position is not straightforward. Computing a motor plan that brings the agent to that posture is even more difficult, since there are multiple possibilities of how the movement may be realized – and the computational complexity increases with the Degrees of Freedom (DoF) of the kinematic chain (e.g., a single arm versus the entire body).

Mainstream theories based on Optimal Control provide a solution to the two inversions that rests upon the optimization of a cost function [2, 3, 4, 5, 6]. However, realizing certain movements such as handwriting or walking is difficult, since not all Bayes-optimal trajectories have well-defined cost functions [7]. Furthermore, the required computations are usually demanding as the model inversion requires considering the effects of the executed actions. Because the latter are perceived with delays that vary across and within sensory modalities[8, 9], they need to be compensated using a forward model, which could introduce additional errors [10].

Active Inference offers an alternative scheme that does not require cost functions [7, 11, 12, 13, 14, 15]. It proposes that agents are endowed with a generative model specifying the dynamics of their hidden states (e.g., hand positions over time) and that desired goals are encoded as priors over the dynamics (e.g., desired hand positions), which act as attracting states. Goal-directed movements are realized by first generating proprioceptive predictions from the hidden states and then minimizing proprioceptive prediction errors, or the discrepancy between predicted and current sensations. Crucially, the mapping between proprioceptive predictions and control signals for the muscles is quite simple and can be implemented with minimal latency using reflex arcs in the spinal cord rather than requiring complex *inverse dynamics* computations [13]. In fact, the inverse model maps from proprioceptive sensations to actions, not from hidden states (either in intrinsic or extrinsic coordinates) to actions, as in Optimal Control [10].

Note that Active Inference only uses reflex arcs as the last stage of control. Complex motor patterns are not constructed at the level of reflex arcs, but rather generated by high-level dynamics that predict specific patterns of proprioceptive predictions [16]. Such predictions encode not just positional terms, but also higher temporal orders [17], allowing the reflex arcs to realize sophisticated instantaneous trajectories that comprise, e.g., velocities and torques [18, 19]. Furthermore, prior expectations over the model dynamics and the entire hierarchy allow immediate first-guess responses, which are eventually refined by subsequent feedback [1].

While there is increasing interest in using Active Inference methods for biological control and robotics [20, 21, 22, 23, 24, 25, 26, 27, 28, 29], their application in realistic settings has been limited so far, given two fundamental issues. First, realizing generative models that afford effective kinematic inversions is challenging. State-of-the-art robotic implementations skip the kinematic inversion and realize movements by only relying on intrinsic coordinates [30, 18]. Despite their effectiveness, such schemes do not allow specifying goals in extrinsic coordinates and hence have several practical limitations. Other approaches embed conventional Optimal Control inversions, such as the Jacobian transpose [31, 10] or the pseudoinverse [32] directly within the dynamics of the hidden states. This has the disadvantage that extrinsic and intrinsic coordinates are mixed up and some computations are duplicated: extrinsic priors are mapped into intrinsic coordinates, which in turn generate the extrinsic predictions to be compared with sensory observations. Hence, the extrinsic generative model has to be embedded also in the dynamics function.

Second, and crucially, the above systems do not model (i.e., maintain probabilistic beliefs about) the entire kinematic chain of the body to be controlled, but only the end effector. Therefore, they fall short of addressing naturalistic bodily movements that require the simultaneous coordination of multiple limbs and joints within a complex kinematic chain – such as the body of living organisms – and they cannot account for movement restrictions due to obstacles.

Here, we introduce a novel Active Inference architecture that resolves the two above limitations and achieves effective and scalable motor control. First, we show that a generative model that maps intrinsic into extrinsic coordinates (IE model) affords an effective kinematic inversion through inference. Second, we show that replicating the scheme of the IE model to mimic the hierarchical structure of the agent’s kinematic chain affords scalable control of bodily movements.

## 2 Results

### 2.1 Kinematic inversion through an Intrinsic-Extrinsic model

Active Inference can afford effective motor control through a kinematic generative model that connects two layers of latent states, encoding intrinsic (***μ***_*i*_) and extrinsic (***μ***_*e*_) coordinates, respectively (see Figure 1A). This “Intrinsic-Extrinsic (IE) model” has two appealing features compared to Optimal Control models and previous Active Inference implementations.

**Figure 1:**
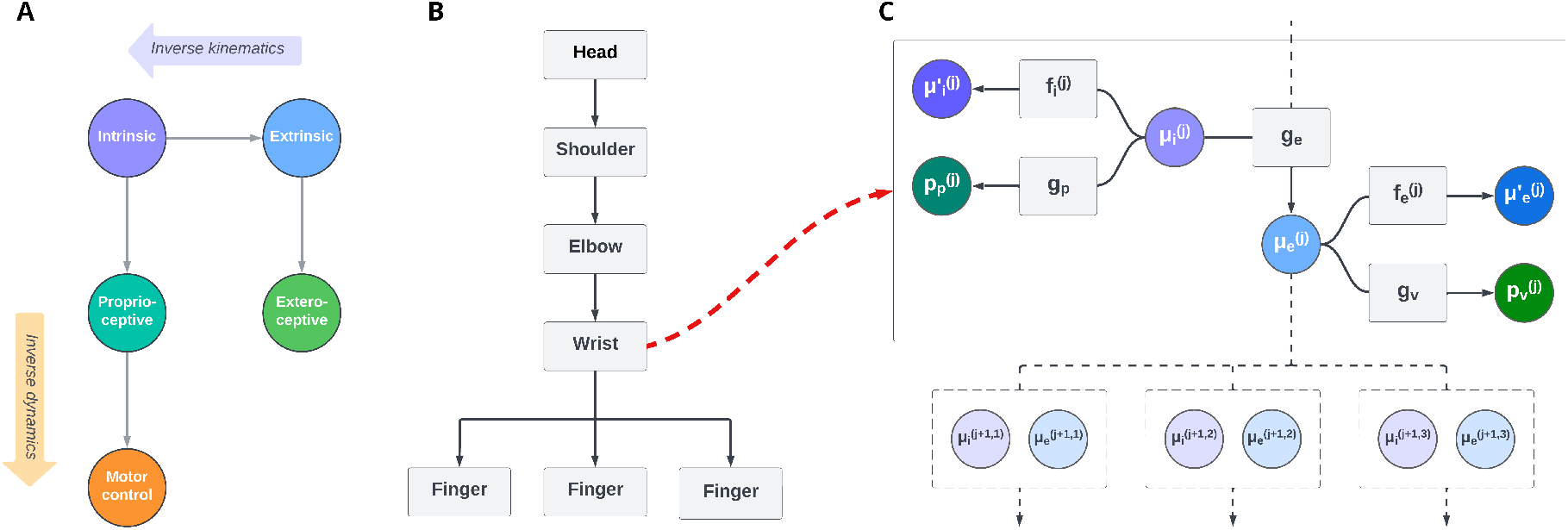
Generative models for deep kinematic inference. (**A**) The Intrinsic-Extrinsic (IE) generative model. The two hidden states are intrinsic and extrinsic coordinates, each with its own dynamics. The proprioceptive generative model extracts joint angles from the intrinsic belief, which are used for both posture inference and movement. At the same time, the exteroceptive model produces a visual prediction (in the simulations reported below, it is approximated by directly using the absolute position of the hand). Inverse kinematics is performed by inference, via backpropagation of extrinsic prediction errors. On the other hand, inverse dynamics from proprioceptive predictions to motor control signals is realized through reflex arcs. (**B**) The deep hierarchical generative model that extends the IE model. The deep model is exemplified for the control of a kinematic plant composed of seven blocks arranged hierarchically, from head to fingers (note that the bottom level is branched, as there are three fingers). Here, the highest (head) level only encodes an extrinsic offset, namely, the initial position and orientation of the origin’s reference frame. (**C**) Factor graph of a single level of the deep hierarchical model. Each block *j* has the same structure as the kinematic generative model shown in Panel A, but with slight differences. In short, intrinsic beliefs 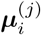 and extrinsic beliefs 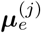 are linked by a generative kinematic model ***g***_*e*_ and they encode information about a single kinematic level. They generate proprioceptive predictions 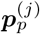 and visual predictions 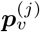 only at their specific level. Intrinsic and extrinsic beliefs have their own dynamics, encoded in the functions 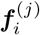 and 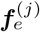, which predict future trajectories and are used for goal-directed behavior. Finally, the extrinsic belief at each level *j* acts as a prior for the level below *j* + 1.

First, the kinematic inversion – or the mapping from extrinsic to intrinsic coordinates – emerges naturally from inference, by inverting the generative mapping from joint angles to Cartesian positions ***μ***_*e*_ = ***g***_*e*_(***μ***_*i*_). This means that the latter does not need to be also specified in the model’s dynamics function [31, 10, 32]. In short, an *extrinsic* attractor (e.g., a point to be reached) first drives the inference of the most likely *intrinsic* hidden state at the level above. Then, the latter affects movement execution through the pathway that generates proprioceptive predictions and finally motor commands.

Second, this scheme allows specifying priors at both intrinsic and extrinsic levels simultaneously, which is needed to realize advanced movements with multiple constraints (e.g., moving the arm while keeping the hand horizontally). Specifying attractors at the intrinsic level is also useful for dealing with particular actions for which this is more natural, or when extrinsic goals are difficult to define. For example, grasping actions can be achieved by specifying priors over object-specific joint configurations (e.g., precision grip for small objects or power grip for large objects).

### 2.2 Extending the IE model to mimic the kinematic chain

The IE model introduced above affords efficient control but still performs the direct kinematics in a single step; in other words, it only predicts the Cartesian coordinates of the end effector (e.g, the hand), not of the entire kinematic chain (e.g., the body). However, the IE model can be extended hierarchically in a straightforward manner, by noting that the direct kinematics of a complex structure can be always decomposed into a sequence of identical transformations (rotations and translations), one for each component of the chain. We exploit this fact to design a hierarchical generative model, composed of as many structurally identical blocks as the number of elements of the kinematic chain where, crucially, each block is an IE model (see Figure 1B-C).

This hierarchical architecture affords direct kinematics by iterating the same operation across all the levels of the kinematic chain, with a biologically plausible *local* message passing. Specifically, the Cartesian position of the tip of the body segment at level *j* is generated from the intrinsic belief over the body segment at the same level and the extrinsic belief over the body segment at the level above *j*− 1. Conversely, the (inverse kinematics) inference of the extrinsic belief at level *j* requires messages from all the extrinsic beliefs at the level below *j* + 1. In fact, as shown in Figure 1B, multiple fingers are attached to the same joint at the bottom level (the wrist). The ramifications can be easily treated by assuming that the inference of the hidden states of the block at level *j* uses the average of the prediction errors generated by all the blocks at the level below *j* + 1 [33]. See Methods for more information.

Overall, this biologically plausible form of local message passing is the same as in hierarchical Predictive Coding for perceptual inference [34, 35]. Here, top-down and bottom-up messages across blocks encode *kinematic predictions* and *kinematic prediction errors*, respectively.

### 2.3 Defining goals for movement using attractive or repulsive forces

In Active Inference, goals for movements can be either imposed as a higher-level prior or embedded into the dynamics function at the same level. The latter approach is usually based on attractors that linearly minimize the distance between the current and desired states. To generate a dynamic goal, the desired state can also depend on some components of the belief itself through a function that embeds movement intentions [36, 37] – see Methods.

The deep kinematic model introduced here permits formulating various kinds of goals at intrinsic and extrinsic levels; for example, the desired angle *θ*^*^ of a joint, the desired absolute position (*x*^*^, *y*^*^), or orientation *ϕ*^*^ of a limb. In addition, it is possible to realize avoidance goals, as shown in Methods. Like attractive forces, repulsive forces are specified for each block of the kinematic chain, which ensures that every segment (rather than just the end effector) is pushed away from an obstacle. Finally, it is possible to define multiple goals simultaneously – such as reaching a particular position with the elbow while maintaining a vertical hand orientation – by combining multiple attractive and/or repulsive forces at different levels. This makes the deep model scalable and versatile, as we show in the next section.

### 2.4 Applications of deep kinematic inference

#### 2.4.1 Reaching task

We demonstrate the capacity of the deep kinematic model in various motor tasks of increasing complexity. We start from the simple reaching task illustrated in Figure 2A, which consists of moving a 4R robotic arm (blue) with realistic joint limits, to reach a static object (red). Here, we compare the performance of the deep kinematic model (*deep*) with the simpler IE model (*IE*) and two alternative controllers based on the Jacobian transpose (*transp*) and the pseudoinverse (*pinv*).

**Figure 2:**
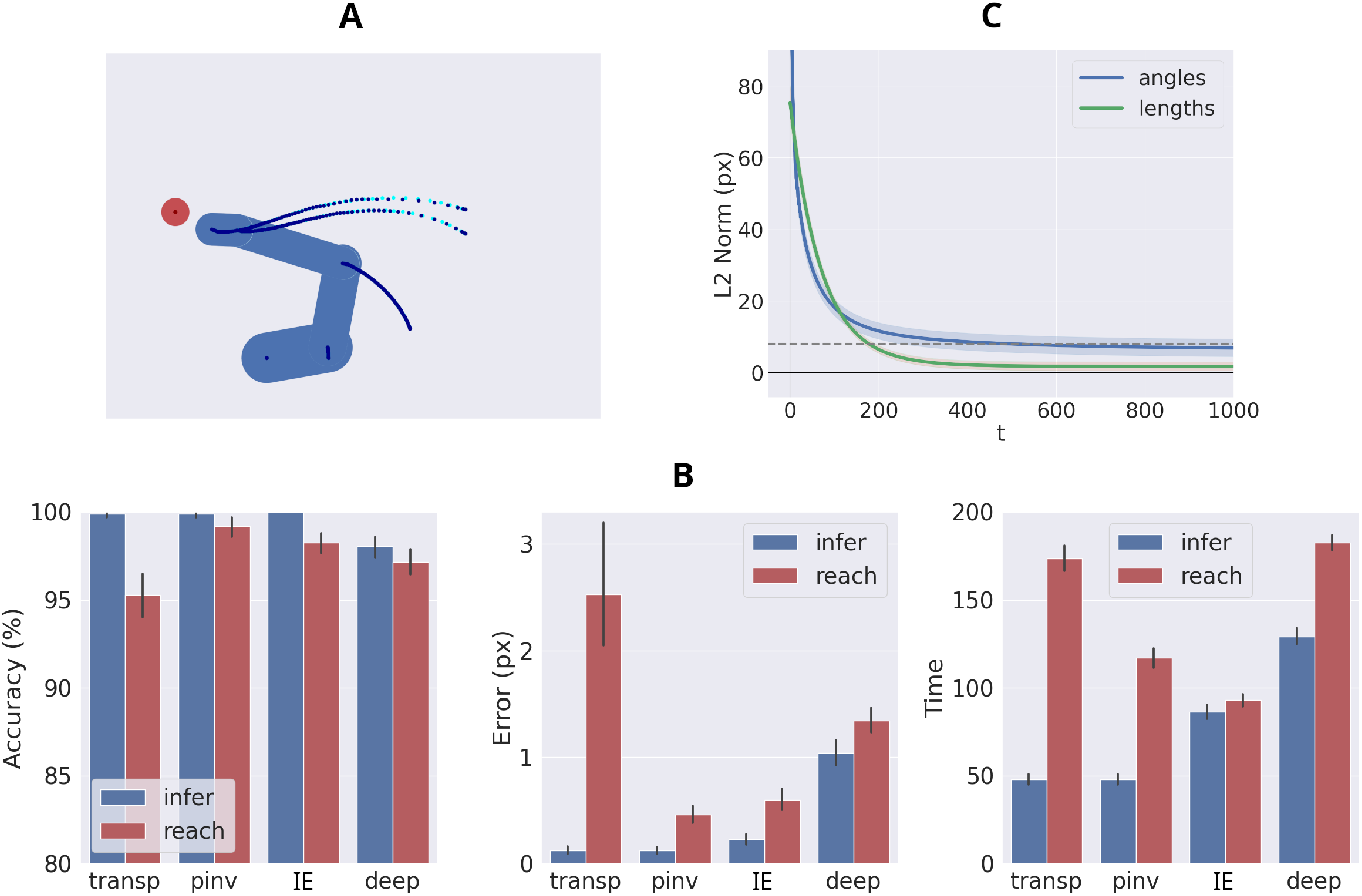
Results. (**A**) Simulation setup: a 4R robotic arm with realistic joint limits reaching a static target (red circle). The trajectory of each joint is shown with blue lines. (**B**) Comparison between the deep kinematic model (*deep*), the simpler IE model (*IE*), and two standard Active Inference controllers based on Jacobian transpose (*transp*) and pseudoinverse (*pinv*). The blue and red bars denote the performance of the models during perceptual inference and reaching, respectively. Note that to better probe the model’s ability to perform perceptual inference, we only allow the model to use visual observations, but not proprioceptive observations during this phase (however, the model can still use proprioceptive observations for reaching). (**C**) Evolution over time of the difference between true and estimated end effector positions (blue line), and between true and estimated lengths of the body segments (red line), aggregated over 1000 trials during inference only. The dashed green line represents the minimum distance defining a successful inference.

The simulation results illustrated in Figure 2B shows the performance of the four models for 1000 trials, with 500 steps for trial. For each trial, the beliefs of the models are initialized with random joint angles and segment positions (a more challenging scenario than starting from congruent intrinsic-extrinsic beliefs). Furthermore, a random target location is sampled and set as the reaching goal for the end effector. We evaluate the performance of the four models with regard to both perceptual inference (blue bars) and reaching ability (red bars), using various metrics (see Figure 2B and Methods).

The performance of the four models is near-optimal during both perceptual inference and reaching. The rare failures are trials in which the models find impossible trajectories and therefore cannot reach the target configurations (note that we did not include any prior over the direction of movement). The performance of the deep model is on par with the other models. The slightly lower performance compared to the IE model is due to the deep model requiring more time to propagate messages across levels, resulting in failed trials when the time steps are limited. The Jacobian pseudoinverse method performs slightly better than the other models, but the transpose method has a much higher final error. Nevertheless, in all cases the average final error is below the minimum distance to consider a trial successful.

### 2.4.2 Learning and adaptation of the kinematic chain

Unlike state-of-the-art implementations, the deep model includes beliefs about the length of the body segments and can infer them over time, based on sensory observations. This may afford the rapid adaptation of the agent to changes in the kinematic chain, e.g., when using a tool that increases the length of the last segment, or when a new joint is added to a specific position. To assess this capability, we performed an experiment in which the beliefs about both angles and body segments are randomly initialized. We evaluated the task over 1000 trials, by comparing the estimated positions of every segment (computed from the joint angles belief) and the estimated lengths with the real values of the same variables. The simulation results illustrated in 2C show that the deep model successfully infers the length of its body segments (green line) in addition to inferring its joint angles (blue line), even within a single trial.

In sum, these results show that the deep model is not less efficient than previous implementations and that different from them, it also affords self-modeling. The next simulations will show that the deep model can be additionally scaled up to account for complex kinematic chains.

#### 2.4.3 Deep inference

Figure 3 shows an evaluation of the deep kinematic model using the same robotic arm as in Figure 2, but equipped with an increasing number of joints, all having equal length. This simulation was assessed on 1000 trials, lasting 2000 time steps for trial. Since we removed the angle limits, the performances are optimal for inference and near-optimal for action. Some trials fail, because the time needed to back-propagate the attractor toward the deepest levels exceeds the time limits, affecting both accuracy and mean error.

**Figure 3:**
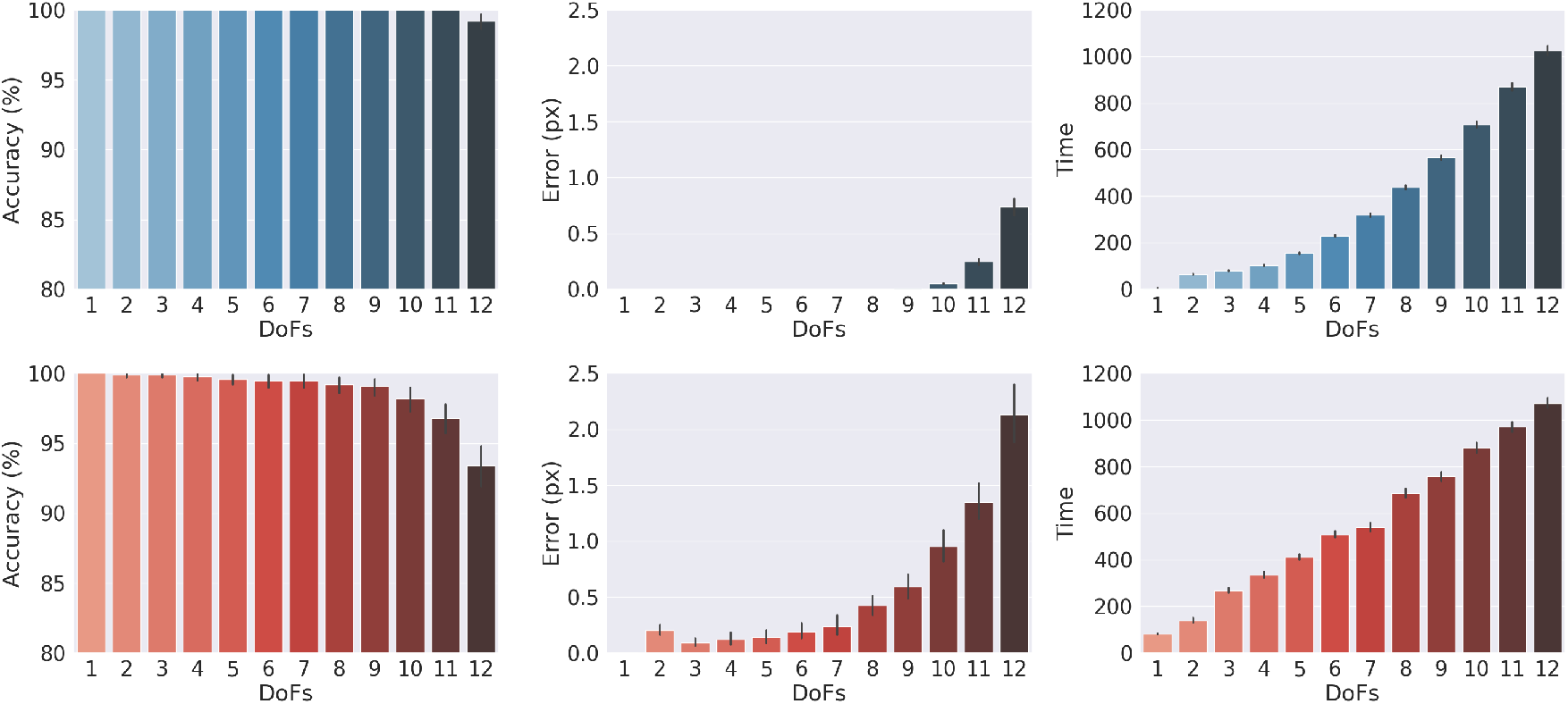
Deep inference. Comparison between hierarchical models with increasing DoF (here, kinematic depth). Inference (top) and reach (bottom) simulations.

### 2.4.4 More complex control tasks

In addition to the previous simulations, the deep model can also control simpler (e.g., arm-like) and more complex (e.g, body-like) structures in a variety of tasks, which require simultaneously tracking a dynamic object (in red) while avoiding another dynamic object (in green) (Figure 4A), making lateral movements while maintaining a vertical orientation of the last segment (Figure 4B), performing circular movements in the Cartesian space (Figure 4C), avoiding a dynamic object with a human-like body (Figure 4D), reaching simultaneously multiple targets with different limbs of a human-like body (Figure 4E), or of a tree-like body with 28 DoF and multiple ramifications (Figure 4F); see the Supplementary Materials for more information about these and various other control tasks.

**Figure 4:**
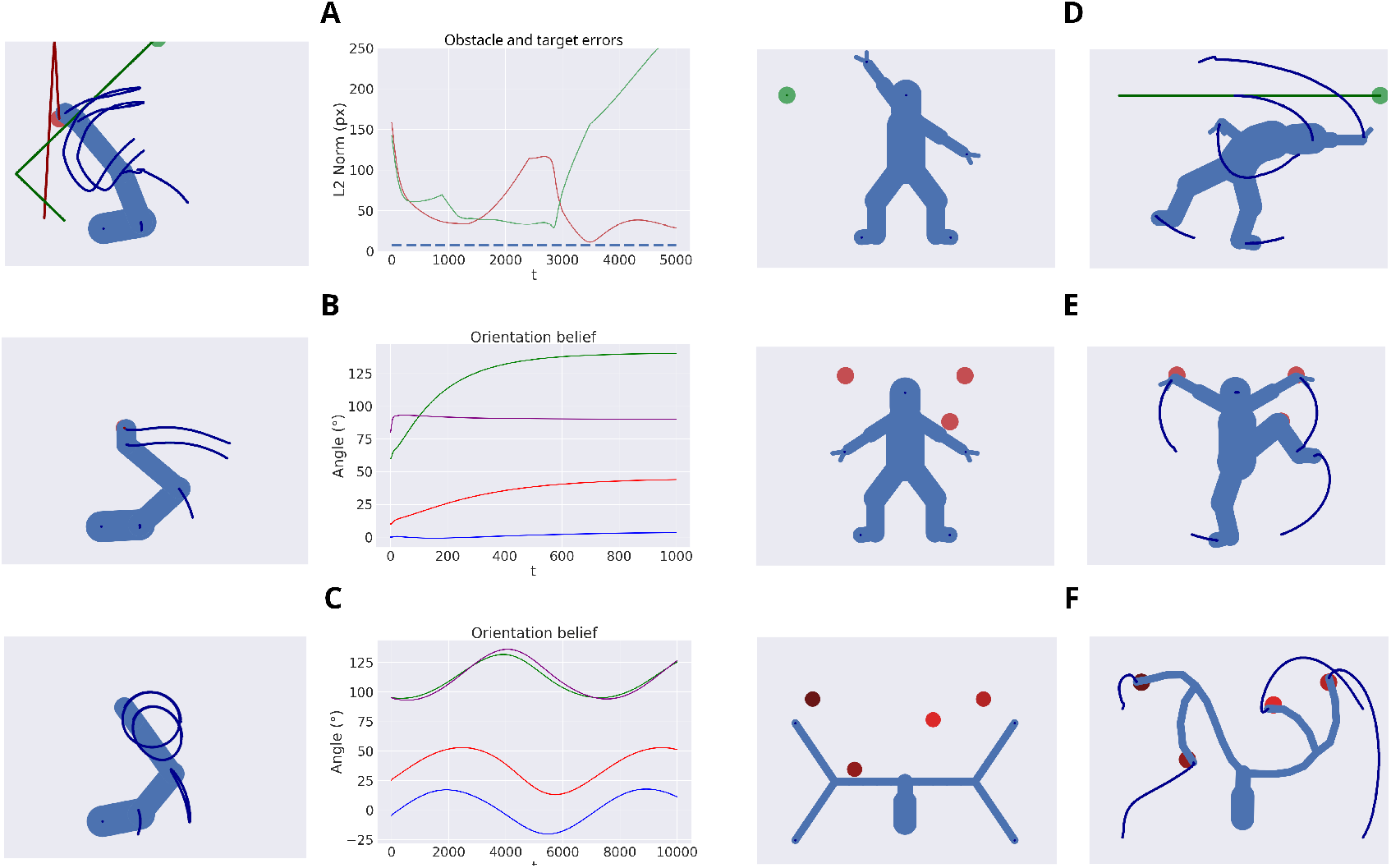
Applications. Track and avoid (**A**), maintain orientation (**B**), perform circular trajectory (**C**), avoid with a full human body (**D**), reach multiple targets with a full human body (**E**), and with a more complex kinematic tree (**F**). The trajectories in the first panel represent the hand-obstacle distance (red line), the hand-target distance (green line), and the minimum distance that the target is considered to be reached (dotted blue line). Instead, the trajectories in the next two panels show the absolute orientation of every joint (in particular, the purple line is the orientation of the hand). More information about the specific tasks and other applications can be found in the Supplementary Materials.

**Figure 5:**
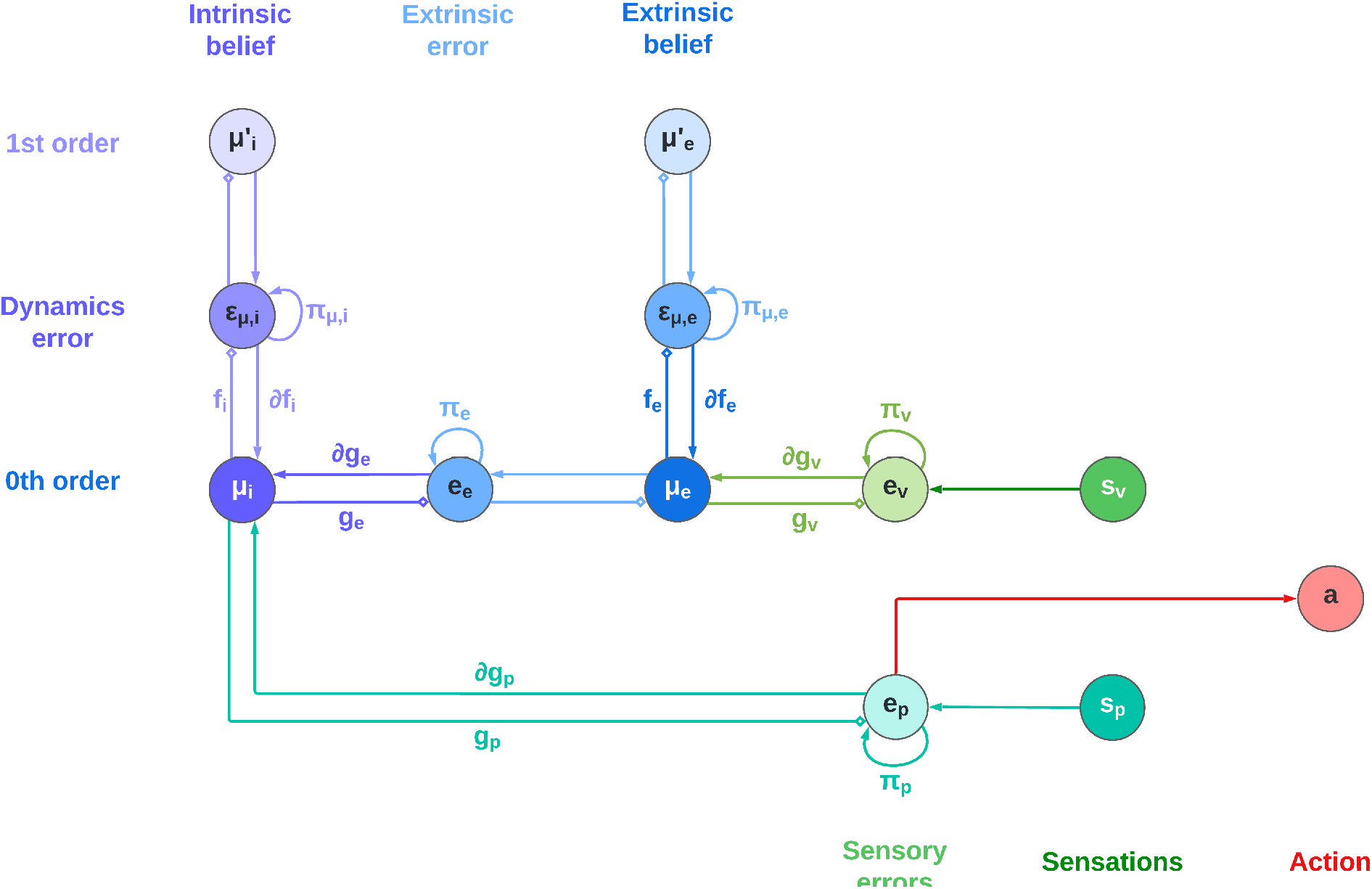
Schematic neural-level implementation of the IE model. The horizontal axis shows the hierarchy between intrinsic beliefs, extrinsic beliefs and sensations. Rather, the vertical axis shows the hierarchy between dynamic levels. The nodes connecting hierarchical levels encode prediction errors. In this schematic, the intrinsic belief directly generates extrinsic predictions of the end effector.

**Figure 6:**
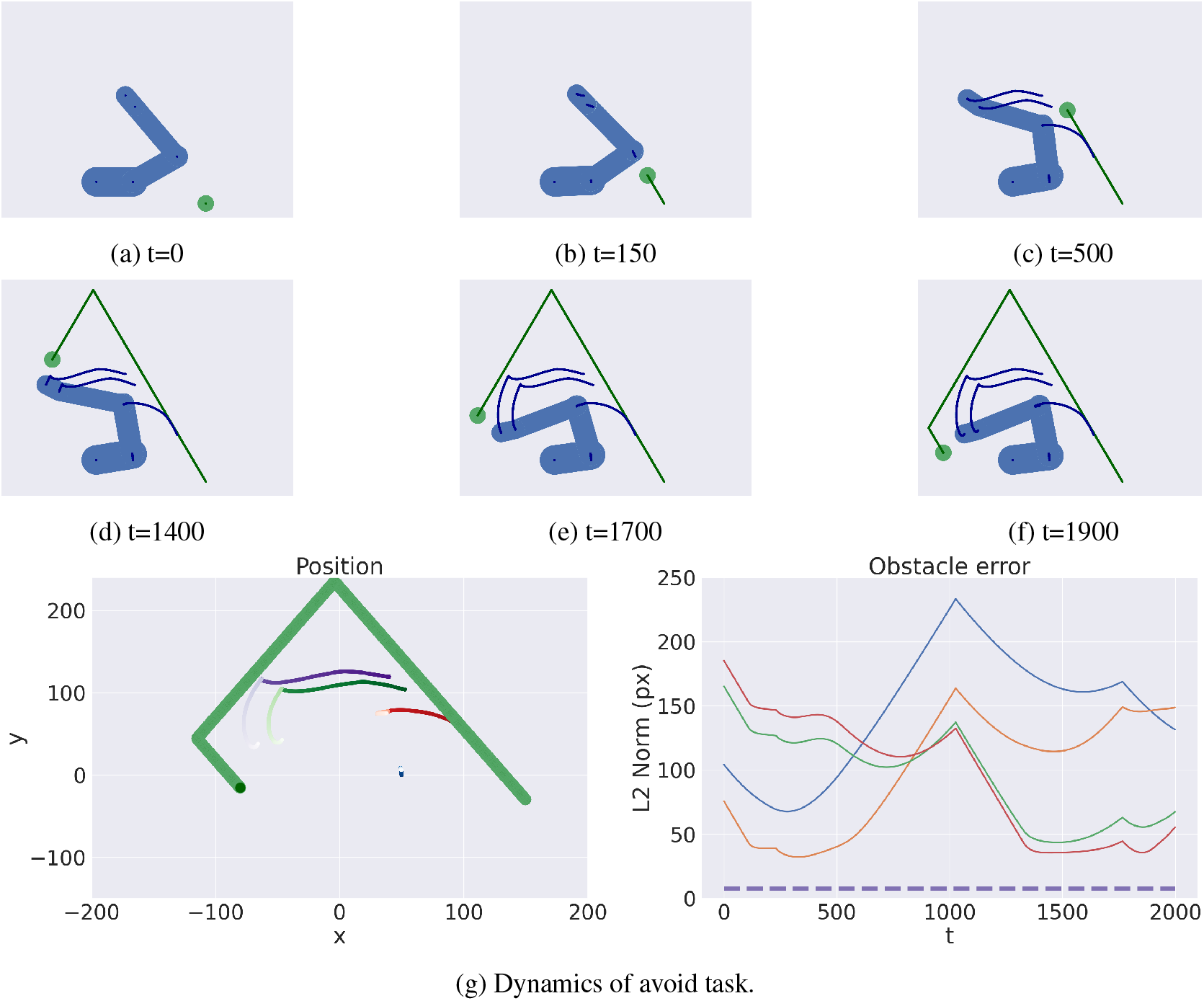
Avoidance of a dynamic obstacle (green circle). (a-f) Sequence of time frames. The arm and obstacle trajectories are shown with blue and green lines, respectively. (g) Trajectory of the arm and the obstacle (left) and distance between the obstacle and each joint (right). The dotted line is the minimum distance required to avoid the obstacle.

**Figure 7:**
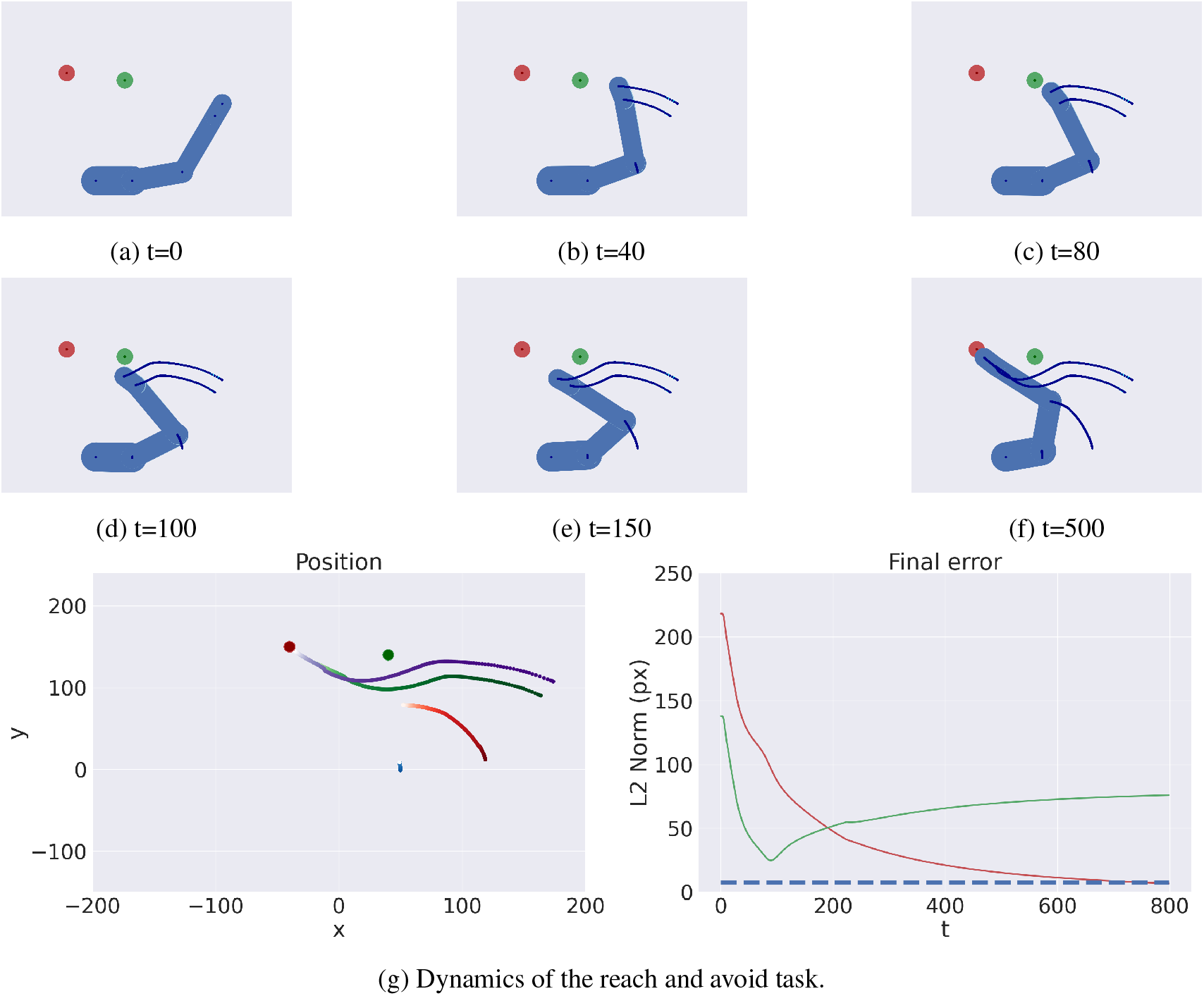
Reach and avoid task. The red and green circles represent the target location and the obstacle, respectively. (a-f) Sequence of time frames. (g) Arm trajectory (left) and distance between the end effector and the target (red line), and between the end effector and the obstacle (green line) The dotted line represents both the reach distance and the minimum distance to avoid the obstacle.

**Figure 8:**
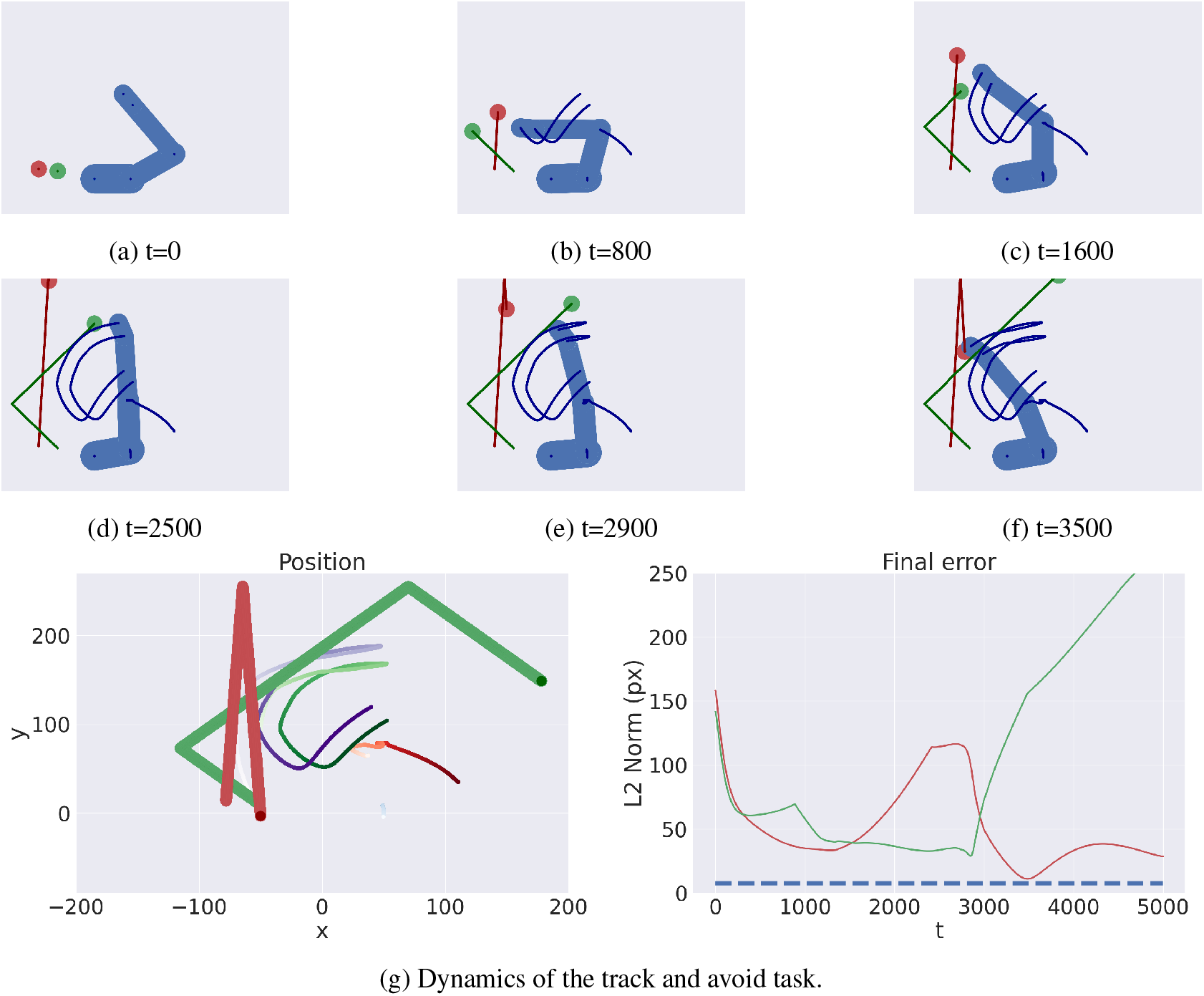
Track and avoid task. (a-f) Sequence of time frames. (g) Arm trajectory (left), distance between the end effector and target (red line), and between the end effector and obstacle (green line). Also see Supplementary Movie 1.

**Figure 9:**
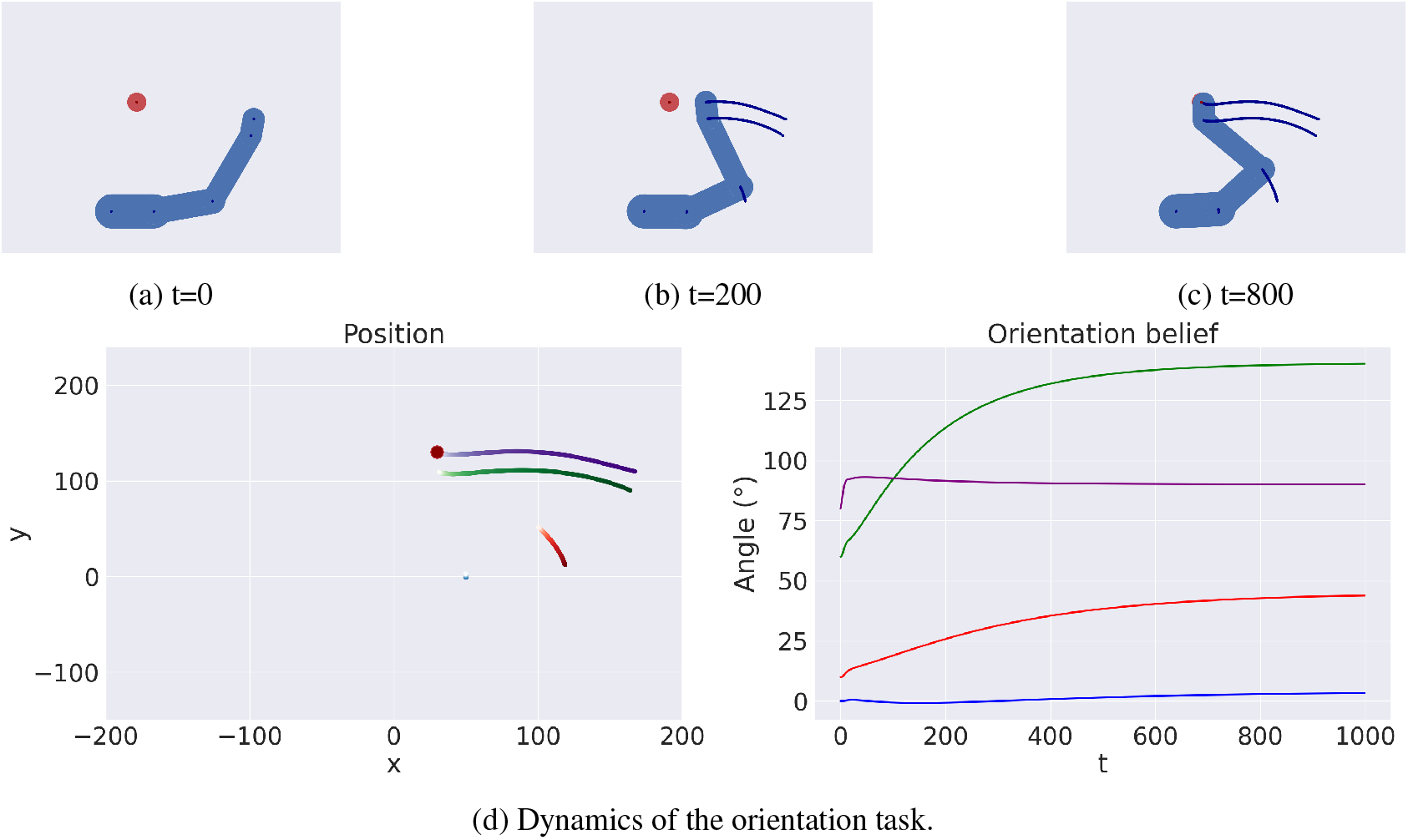
Maintain the same orientation. (a-c) Sequence of time frames. (d) Arm trajectory (left) and absolute orientation of every joint over time (right). The last joint is represented in purple. Also see Supplementary Movie 2.

**Figure 10:**
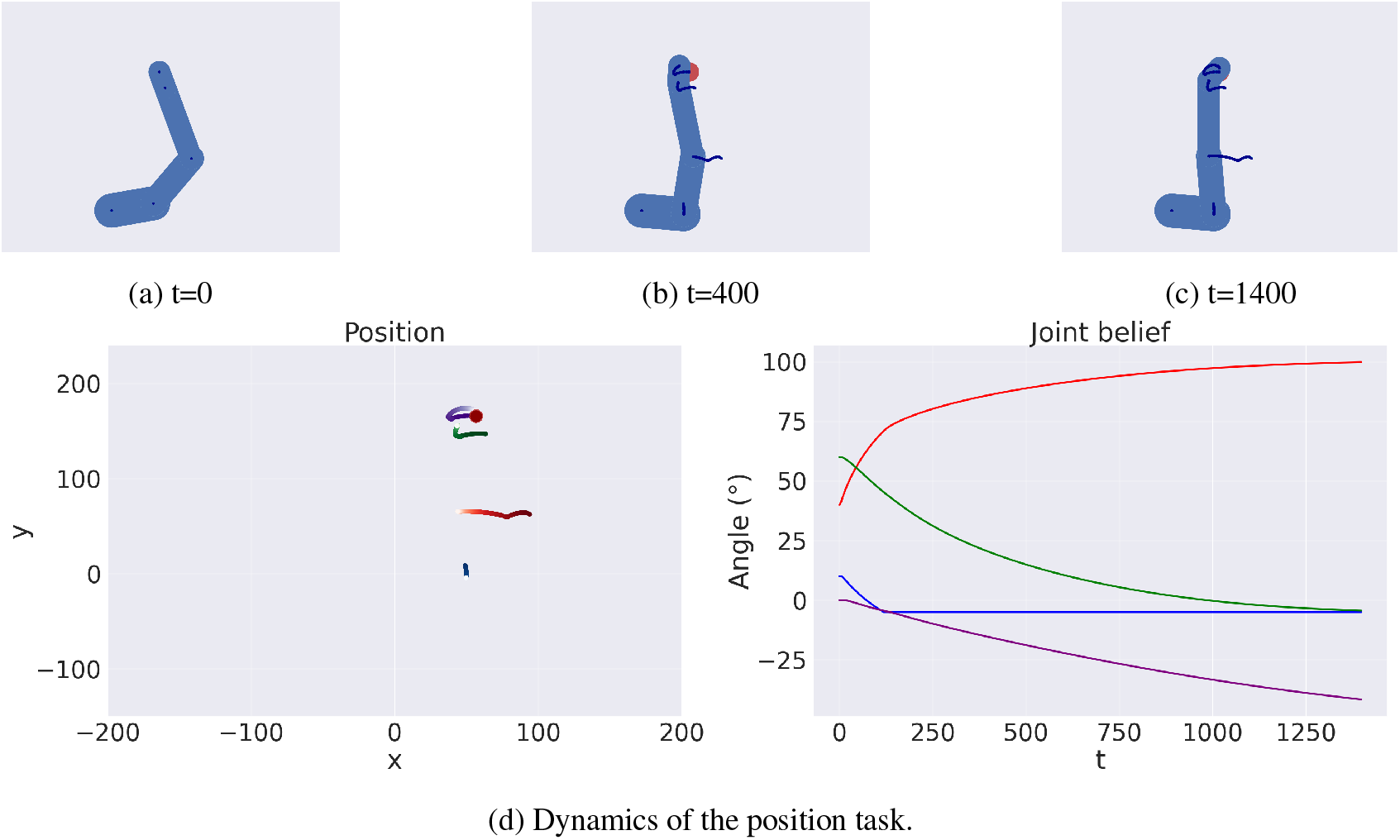
Maintain the same position. (a-c) Sequence of time frames. (d) Arm trajectory (left) and relative angles of every joint over time (right). The shoulder joint is represented in red.

**Figure 11:**
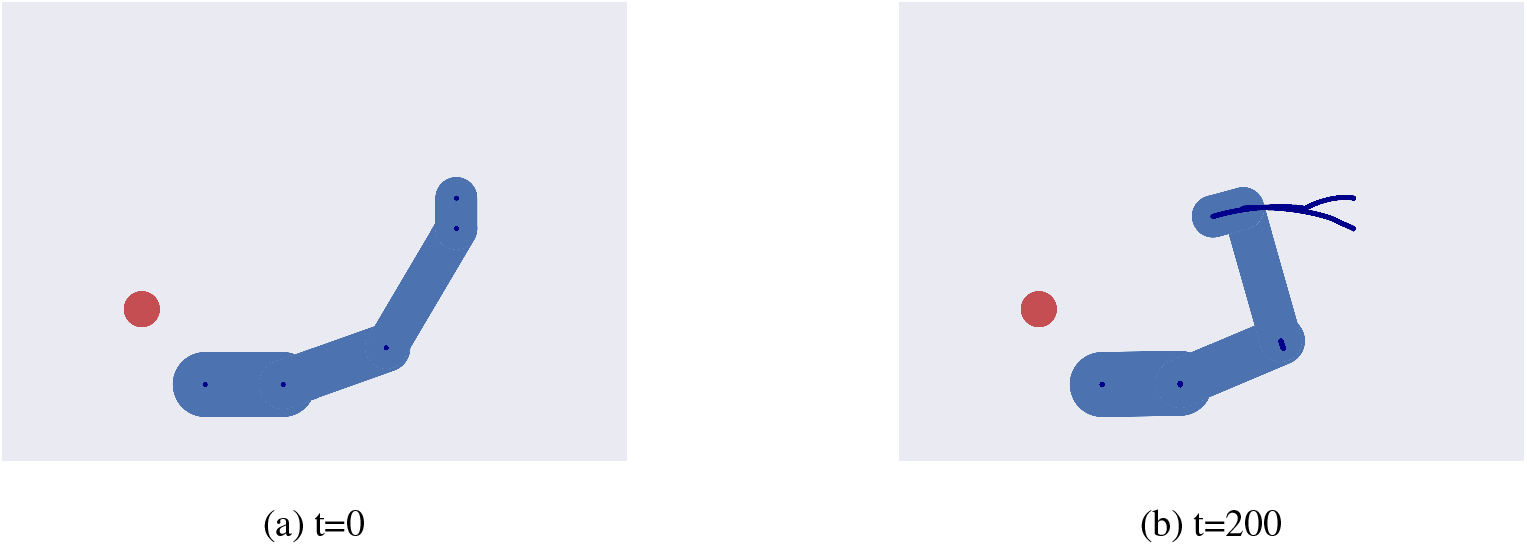
Target pointing. Although the goal is only specified at the level of the last joint, the agent will move freely, until the end effector points towards the target.

**Figure 12:**
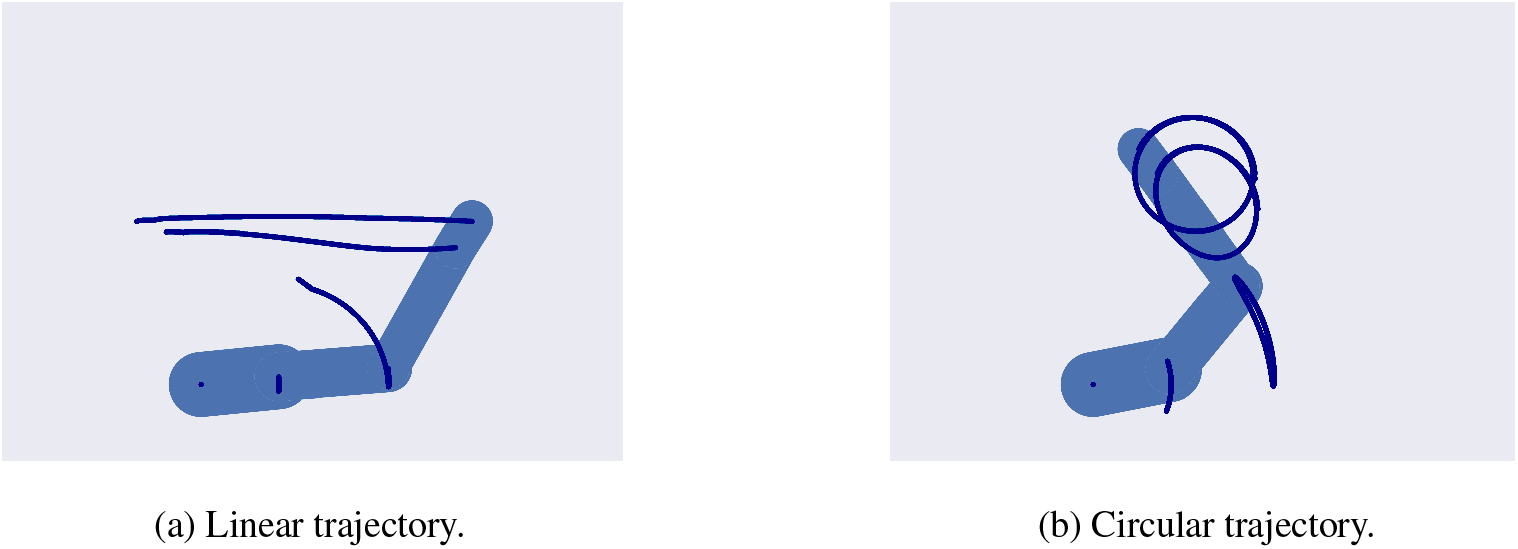
Trajectory planning in the Cartesian domain. Slowing down sufficiently the extrinsic dynamics allows to perform linear (left) and circular (right) trajectories. Also see Supplementary Movie 3.

**Figure 13:**
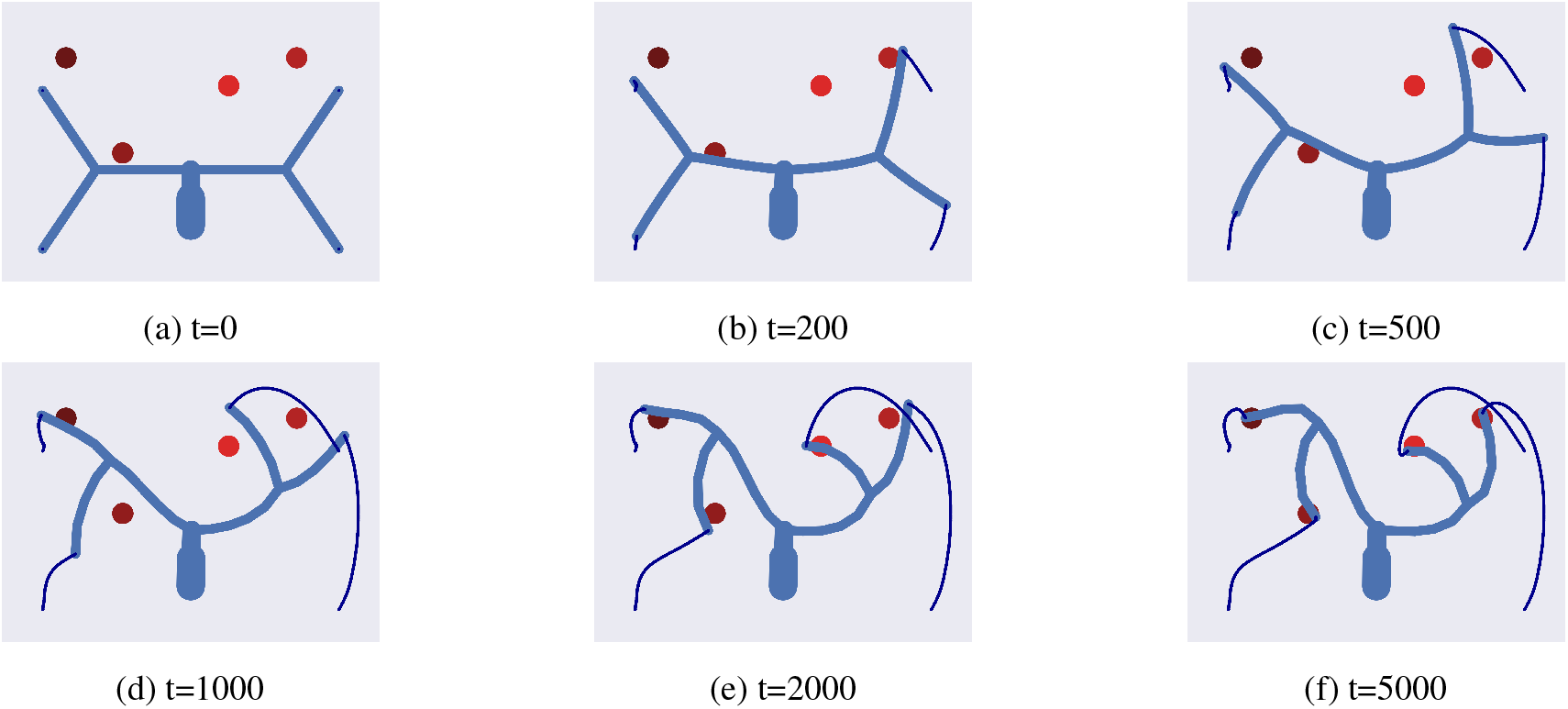
Controlling complex Kinematic Structures. The goal consists in reaching 4 different target locations with a 28-DoF agent. Also see Supplementary Movie 4.

Crucially, the same deep model can achieve all these (and other) tasks by simply specifying different goals, using attractive or repulsive forces, or a combination of both, without having to define ad-hoc cost functions. Compared to other approaches, the deep model is therefore particularly easy to scale up to address complex kinematic structures and control problems with multiple goals.

## 3 Discussion

Active Inference proposes that goal-directed movements are realized by producing predictions with a generative model of the coupled agent-environment system, and then minimizing the errors between predicted and current sensations. Despite its strong theoretical appeal [1], this framework has enjoyed limited applications in motor control and robotics, given the challenge of specifying appropriate generative models to map different (extrinsic and intrinsic) coordinates of movement, and to control complex kinematic structures such as the human body.

Here, we show that a generative (IE) model that maintains distinct beliefs over intrinsic and extrinsic coordinates affords effective control of goal-directed movements, because it allows defining both intrinsic and extrinsic goals as attractive (or repulsive) states and solving inverse kinematics via (active) inference, which is typically challenging in other frames like Optimal Control. Furthermore, we show that a deep hierarchical extension of the IE scheme drastically improves its scalability. In particular, replicating the same IE generative model for each block of the kinematic chain simplifies the computations of the direct kinematics and permits controlling complex kinematic structures that require the simultaneous coordination of multiple segments.

The proposed deep hierarchical architecture has four main advantages compared to Optimal Control schemes or state-of-the-art models in the Active Inference literature. First, it is efficient: performing the kinematic inversion through inference drastically reduces its computational cost. Second, it is scalable: it is possible to design generative models for sophisticated kinematic chains by simply connecting multiple IE models. Since the overall generative model always mimics the entire kinematic chain, adding or removing new segments (e.g., to model tool use) simply requires adding or removing the corresponding blocks. Third, it affords self-modeling [38] by continuously inferring the kinematic structure online and rapidly re-adapting to changes in limb lengths. Fourth, it affords a biologically plausible form of inference that only uses *local* message passing and *asynchronous* computations, akin to hierarchical Predictive Coding for perception [34, 35, 11, 39].

The behavior of the deep kinematic model, which replicates the same computational block across various levels, bears some analogies with deep neural networks. It is common to use the latter (e.g., Variational Autoencoders) as generative models for Active Inference agents [30, 36]. Despite their effectiveness, the deep networks are treated as black boxes during the free energy minimization: the belief over hidden states only receives a single gradient and it is unaware of the internal computations performed by the backpropagation algorithm. This has the downside that the weights of the generative model have to be learned a priori and remain constant throughout the Active Inference process – hence, the agent cannot adapt them when it receives novel sensory observations. Furthermore, the agent has no control over the dynamics of the generative model and cannot control the intermediate levels of the deep network toward potentially preferred states. In other words, almost all the hard work is delegated to the deep network and the only job left to the Active Inference agent is inferring the highest level of the hidden states. In contrast, one appeal of Predictive Coding methods is that they can simultaneously perform hidden state inference and learning [34, 35, 11, 40]. Designing a hierarchical structure that uses the same rules of free energy (or prediction error) minimization throughout the levels means that the agent can constantly modify its internal models to match both sensory observations and prior expectations.

In this study, we focused on the theoretical aspects of controlling a complex hierarchical structure with multiple constraints. For this reason, we used a relatively simple velocity-controlled system, without considering higher temporal orders. This implies that the dynamics functions illustrated lack some of the constraints required to smoothly actuate a real system. However, such constraints can be easily included in the Active Inference scheme used here, by imposing priors at specific levels of the hierarchy. We provided one example in the Results section, when we discussed how the system can incorporate specific affordances, but other useful examples can be made. Given the simple analytical form of the kinematic model, one can achieve movements with the desired smoothness by expanding the generalized state space and defining dynamics functions of increasing temporal orders as we did for our velocity-controlled scheme. As a result, jerk minimization can be realized by maintaining an intrinsic belief with temporal orders up to the 4th and imposing a limit at the last level [18]. Similarly, an attractor could be defined at the 3rd level, setting specific torques to a joint and allowing one to simulate advanced limb dynamics [19]. Further, singularities can be avoided by predicting their influence in advance and embedding them at the intrinsic level [12]. Last, advanced movements beyond reaching and avoiding tasks – such as those requiring high-level planning – may be achieved through a hybrid scheme, in which a discrete model plans an optimal sequence of actions that are in turn realized by a continuous model like the one exemplified in this study [17]. In other words, the hybrid scheme allows the agent to model the environment at a more abstract level, realizing multi-step reaching movements [41], object manipulation tasks, and plan trajectories that avoid the local minima that arise with Artificial Field Potentials. We plan to simulate these capabilities of hybrid Active Inference in future works.

Finally, although we exemplified a deep kinematic inference for the control of human body movements, the same design could be used to control other, potentially more sophisticated structures, composed of multiple ramifications. The deep kinematic inference approach can therefore pave the way for the efficient and biologically plausible control of general actuated systems.

## 4 Methods

### 4.1 Hierarchical Active Inference

The core of Active Inference is the use of a generative model to generate predictions and minimize any prediction error resulting from discrepancies between predictions and observations. The generative model depends on three elements encoded in generalized coordinates of increasing temporal orders (e.g., position, velocity, acceleration, etc.): hidden states 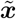, hidden causes 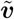 and sensory signals 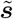. These elements are related by a nonlinear system specifying the generation of sensory signals and the evolution of latent states over time:

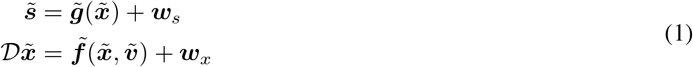

where 𝒟 is a differential operator that shifts all the temporal orders by one, i.e.: 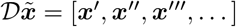, while ***w***_*s*_ and ***w***_*x*_ are noise terms assumed to be sampled from a Gaussian distribution. The associated joint probability is factored into independent distributions:

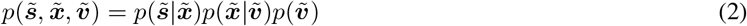

where each distribution is usually approximated by Gaussian functions:

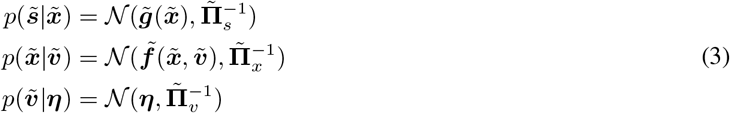

expressed in terms of precisions or inverse variances 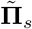, 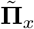, and 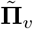. Following a variational inference approach [42], these distributions are inferred through approximate posteriors 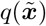 and 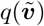. Minimizing the Variational Free Energy (VFE) ℱ, defined as the difference between the KL divergence of the (real and approximate) posteriors and the log evidence:

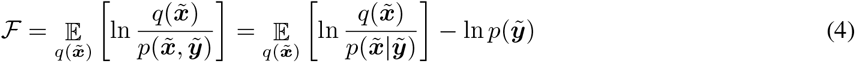

reduces to the minimization of prediction errors, and the updates of the beliefs 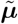 and 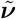 respectively over the hidden states and hidden causes become:

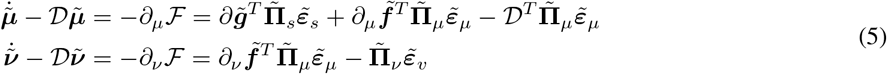

where 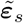, 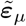, and 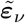 are respectively the prediction errors of sensory signals, dynamics, and priors:

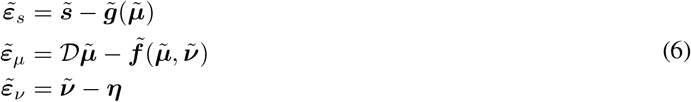

A simple formulation of Active Inference can deal with several tasks, but its main strength relies on a hierarchical structure that allows the brain to learn the inherently hierarchical relations between sensory observations and their causes [17].

This structure can be easily scaled up by linking every hidden cause with another generative model; thus, the prior becomes the prediction from the layer above, while the observation becomes the likelihood of the layer below:

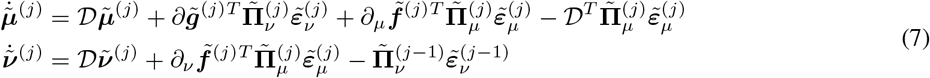

where:

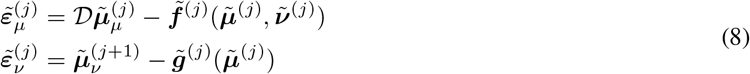

On the other hand, action is realized by minimizing the proprioceptive component of the VFE with respect to the motor control signals ***a***:

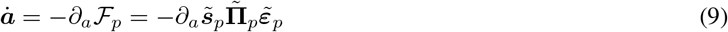

where *∂*_*a*_***s***_*p*_ is the partial derivative of proprioceptive observations with respect to the motor control signals, 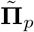 are the precisions of proprioceptive generative models, and 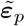 are the generalized proprioceptive prediction errors:

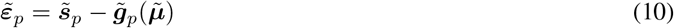

### 4.2 Kinematic models: belief update

In the IE model (shown in Supplementary Figure S1), the intrinsic belief encodes every joint angle in parallel while the extrinsic belief only encodes the Cartesian position of the end effector. For a 2R robotic arm, such beliefs are related by:

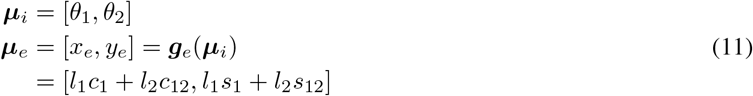

where ***g***_*e*_ is a generative model performing forward kinematics, and we used a compact notation to indicate the sine and cosine of the angles:

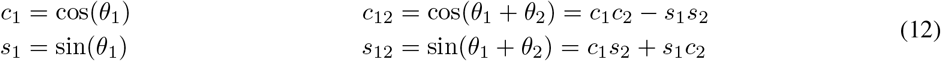

The update equation for the intrinsic belief is then a combination of proprioceptive likelihood, extrinsic likelihood, and intrinsic attractor dynamics:

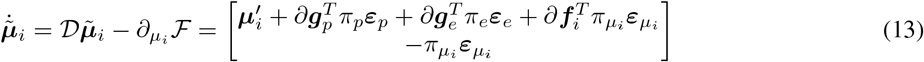

The form of the dynamics function ***f***_*i*_ is explained in the following sections. The kinematic inversion is performed through the gradient *∂****g***_*e*_:

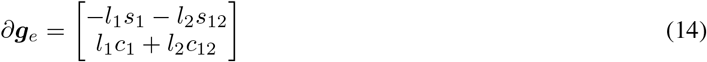

Similarly, the update for the extrinsic belief is a combination of intrinsic prior, visual likelihood, and extrinsic attractor dynamics:

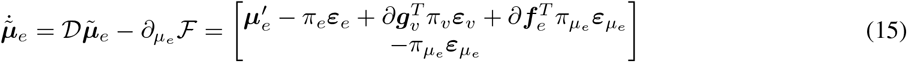

Concerning the deep kinematic model, if we consider a simple 2D arm, each block is composed of an intrinsic belief over pairs of joint angles and segment lengths 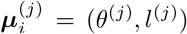 and an extrinsic belief over the position of the segment’s extremity and its absolute orientation 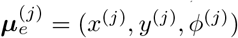. The resulting kinematic generative model is:

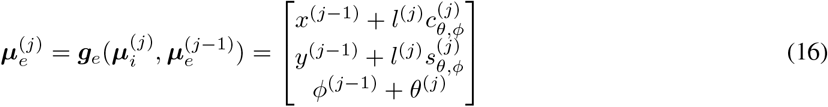

The update of the intrinsic belief is equivalent to the IE model:

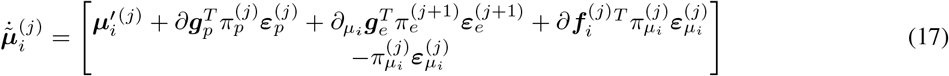

However, the gradient of the kinematic generative model with respect to the intrinsic belief is simpler, since it only depends on the intrinsic components of that level:

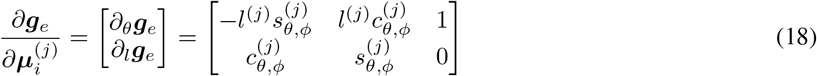

Note that the second row of the gradient in Equation 18 allows inferring and learning segment lengths.

The update of the extrinsic belief still includes every component of the IE model, but with the addition of the extrinsic likelihood from the next layer *∂*_*e*_***g***_*e*_, which in this case is the sum of all the segments it is attached to:

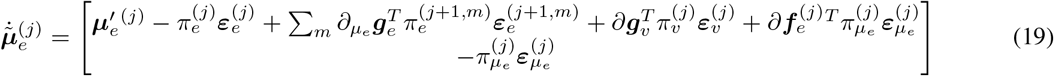

The gradient of this new term connects every layer in the hierarchy since intrinsic beliefs communicate with their extrinsic predictions by:

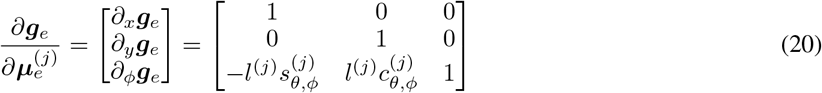

### 4.3 Ramifications

If we consider a ramification with *M* extrinsic outputs 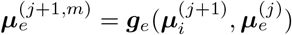, the update equation for the parent node 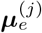 is proportional to:

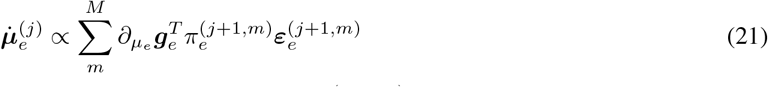

where 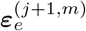 is the extrinsic prediction error of block (*j* + 1, *m*) and 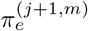 is the corresponding precision. The latter thus weighs the contribution for each branch of the level.

### 4.4 Defining attractive goals and repulsive forces

Goals can be flexibly formulated at both intrinsic and extrinsic levels through an intention function ***i***(***μ***) that linearly combines the current belief with biases ***h***^*^ to define the desired future state ***μ***^*^:

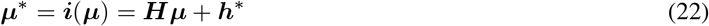

In brief, the matrix ***H*** performs a transformation of the current belief: manipulating and combining only specific components affords a dynamic behavior (e.g., reaching a moving target), while using an identity transformation for the other components lets them free to change since their prediction error would be zero. Instead, the vector ***h***^*^ is used to impose a static configuration.

This function could specify, for example, the desired angle *θ*^*^ of a joint (i.e., an intrinsic goal), the desired absolute position (*x*^*^, *y*^*^) or the orientation *ϕ*^*^ of a limb (i.e., extrinsic goals):

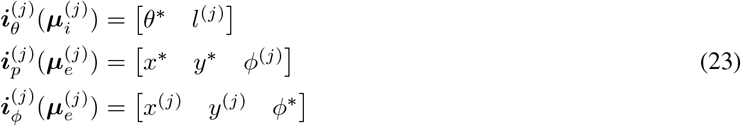

Attractors then linearly minimize the distance between the current and desired states, i.e., ***f***_*a*_(***μ***) = *k*_*a*_***e***_*a*_, where *k*_*a*_ is an attractive gain and ***e***_*a*_ = ***μ*** *−* ***μ***^*^.

In turn, following the Artificial Field Potentials theory, repulsive forces can be defined as:

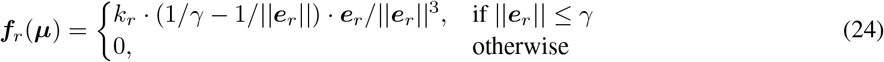

where *k*_*r*_ is a repulsive gain, *γ* is the range of influence of the repulsive force, and ***e***_*r*_ = ***μ μ***^#^ is the difference between the belief and a repulsive state ***μ***^#^ defined flexibly in the same way as the goal state ***μ***^*^. At this point, one can combine attractive and repulsive forces into the dynamics function of a single level *j* in order to achieve a complex behavior – e.g., reaching a target with the end effector while avoiding an obstacle.

### 4.5 Assessment Metrics

The three metrics used to evaluate perceptual inference are: (i) *perception accuracy*: success in finding a joint configuration corresponding to the true target location within 8 pixels; (ii) *perception error*: *L*^2^ distance between the true and estimated target positions at the end of the trial; (iii) *perception time*: number of time steps needed to successfully estimate the target position.

The three metrics used to evaluate reaching are (iv) *reach accuracy*: success in approaching the target within 8 pixels; (v) *reach error*: *L*^2^ end effector-target distance at the end of the trial; (vi) *reach time*: number of time steps needed for the end effector to reach the target in successful trials. Note that the reaching task is more challenging than the inference task, since it requires inferring the final arm configuration based on information about only the position of the last segment.

## 5 Acknowledgments

This research received funding from the European Union’s Horizon H2020-EIC-FETPROACT-2019 Programme for Research and Innovation under Grant Agreement 951910 to IPS, the European Union’s Horizon 2020 Framework Programme for Research and Innovation under Grant Agreement 945539 to GP, the European Research Council under the Grant Agreement 820213 to GP, and the Italian Ministry for Research MIUR, under Grant Agreement PRIN 2017KZNZLN to IS. The funders had no role in study design, data collection and analysis, decision to publish, or preparation of the manuscript.

## 6 Supplementary Materials

### 6.1 Applications

In this section, we show further practical applications of the deep kinematic model.

#### 6.1.1 Avoid task

In the avoid task, the agent has to avoid a moving obstacle with every joint, see Supplementary Fig. 6. In this task, the agent is only subject to repulsive potentials, specified at every level of the hierarchy. At trial onset (t=0), the obstacle starts in the lower right corner with a random direction. At t=150, the agent starts perceiving a repulsive force and moves to avoid the obstacle with the elbow. At t=500, the agent successfully avoids the obstacle. At t=1400, as the obstacle approaches again the arm, the agent perceives another repulsive potential, but now at the end effector position, and avoids again the obstacle in 300 time steps. Finally, at t=1900, the agent performs another slight adjustment. In all cases, it manages to successfully keep at a safe distance, as shown in the right panel of Supplementary Fig. 6g.

#### 6.1.2 Reach and avoid task

In the reach and avoid task, the agent has to reach a target location with the end effector, while avoiding an obstacle located in the middle of the trajectory, see Supplementary Fig. 7. Here, the agent is subject to both attractive potentials (only for the end effector) and repulsive potentials (for every joint), encoded as extrinsic beliefs. At t=0, both target and obstacle spawn at random positions. At t=40, the agent moves only by the attractive force. At t=80, the agent begins perceiving the obstacle and curves its trajectory. At t=150, it has successfully avoided the obstacle, and it moves again only by the attractive potential - although it may still be influenced by repulsive potentials from intermediate joints.

#### 6.1.3 Track and avoid task

The track and avoid task, which is similar to the previous one, except that the target moves, see Supplementary Fig. 8. The goal of the agent is to track the target for the whole trial, while avoiding the dynamic obstacle. At t=0, the target and the obstacle spawn in close positions but with different directions. The agent starts to track the target, but at t=1600 it perceives a repulsive force that is stronger than the attractive force from the target. At t=2900, it manages to avoid the obstacle and continues tracking the target.

#### 6.1.4 Maintain final orientation task

In this task, the agent has to reach a target location while maintaining a vertical end effector orientation during the whole trajectory (as if it had a glass full of water in the arm and did not want the water to fall), see Supplementary Fig. 9. In this task, the agent has two goals. The former consists in setting the end effector position to the target location, as in the previous tasks. The latter sets the end effector’s absolute orientation to 90 degrees. Different from the previous tasks, we thus specify every component of the extrinsic belief of the last layer:

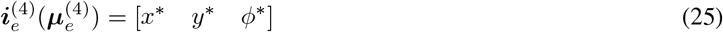

As shown in Supplementary Fig. 9d, the agent achieves the task successfully, by correctly maintaining the vertical end effector orientation while moving from the initial to the final location.

#### 6.1.5 Maintain final position task

In this task, the agent has to set the relative joint angle of the shoulder to a specific orientation (here, 100 degrees) while maintaining a fixed end effector position, see Supplementary Fig. 10. This task requires fulfilling two goals, which are specified at different levels and reference frames. The goal to set the shoulder joint to a specific angle is defined in the intrinsic reference frame, while the goal to keep the end effector at a fixed target position is defined in the extrinsic domain.

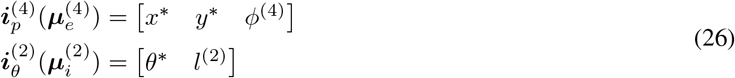

As shown in Supplementary Fig. 10d, the agent reaches the desired shoulder angle correctly and rotates the wrist to maintain the same position; yet it slightly moves the end effector during the trajectory, since the joint limits do not allow perfect tracking.

#### 6.1.6 Target pointing task

Sometimes we want to point to a target, rather than reach it (see Supplementary Fig. 11). While a reaching goal is inappropriate, it is possible to impose an absolute orientation over a segment equal to the direction of the vector from the segment’s extremity to the target:

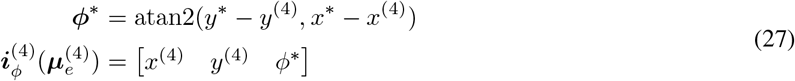

Also here the inference quickly drives the agent to implement the desired movement (Supplementary Fig. 11b).

#### 6.1.7 Trajectory planning in Cartesian space

Defining attractors in the extrinsic belief is also useful for realizing optimal trajectories in the Cartesian space, as it is possible to define every sort of path, velocity, and acceleration profiles. However, since the extrinsic prediction errors are backpropagated through the inverse kinematic model, the actual movement does not necessarily result in the desired trajectory. Imposing a prior of a linear velocity over the 1st-order extrinsic dynamics results in a slightly curved movement, if the update dynamics of both beliefs are similar. However, if the extrinsic update is sufficiently slower, the intrinsic belief will follow the desired trajectory, thus allowing to perform linear or circular movements, as shown in Supplementary Fig. 12.

#### 6.1.8 Controlling complex kinematic structures

The deep model can deal with complex kinematic structures, such as the one shown in Supplementary Fig. 13, composed of 28-DoF and linked in a way as to form a tree of depth 3. The goal of the agent is to reach 4 different target locations with the end of each branch. This task is more difficult than the previous ones: apart from the much higher number of joints, the inference for the high-level layers should balance the contributions from different branches. Yet, the deep model is able to complete the task successfully, despite requiring a considerably higher number of time steps.

Similarly, it is possible to simulate human full-body kinematics. To exemplify this, we performed two tasks, which required controlling a simplified human body composed of 23-DoF, to reach 3 different target locations with the left knee and both hands (Fig. 14) and to avoid a moving obstacle with the whole body (Fig. 15). In both simulations, the remaining segments freely move - depending on which ones are constrained - by backpropagation of errors throughout the whole kinematic chain, thus realizing realistic body movements. Other intentions can be imposed if the goal is only to move the desired joints.

**Figure 14:**
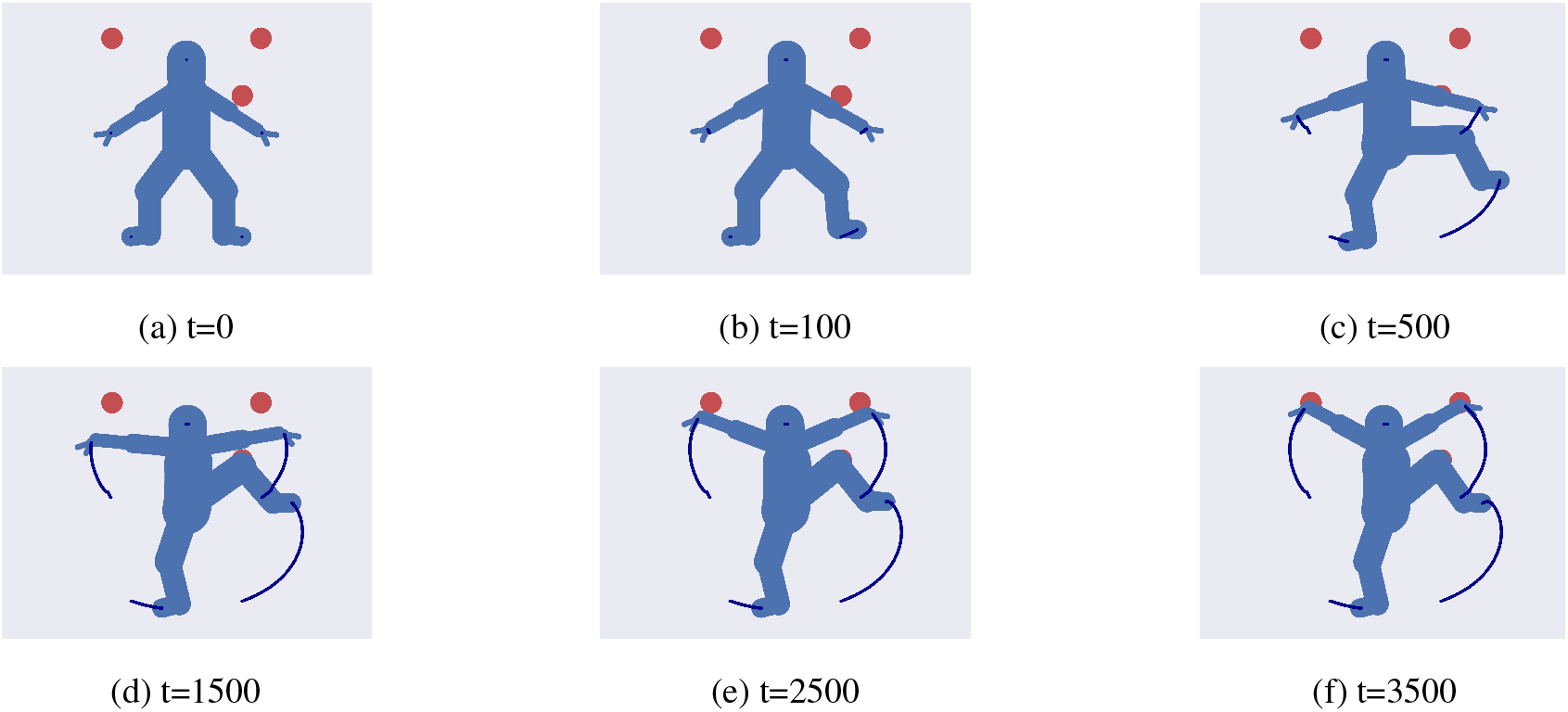
Controlling a simplified human body composed of 23-DoF. The goal is to reach 3 different target locations, with the left knee and the two arms. Also see Supplementary Movie 5.

**Figure 15:**
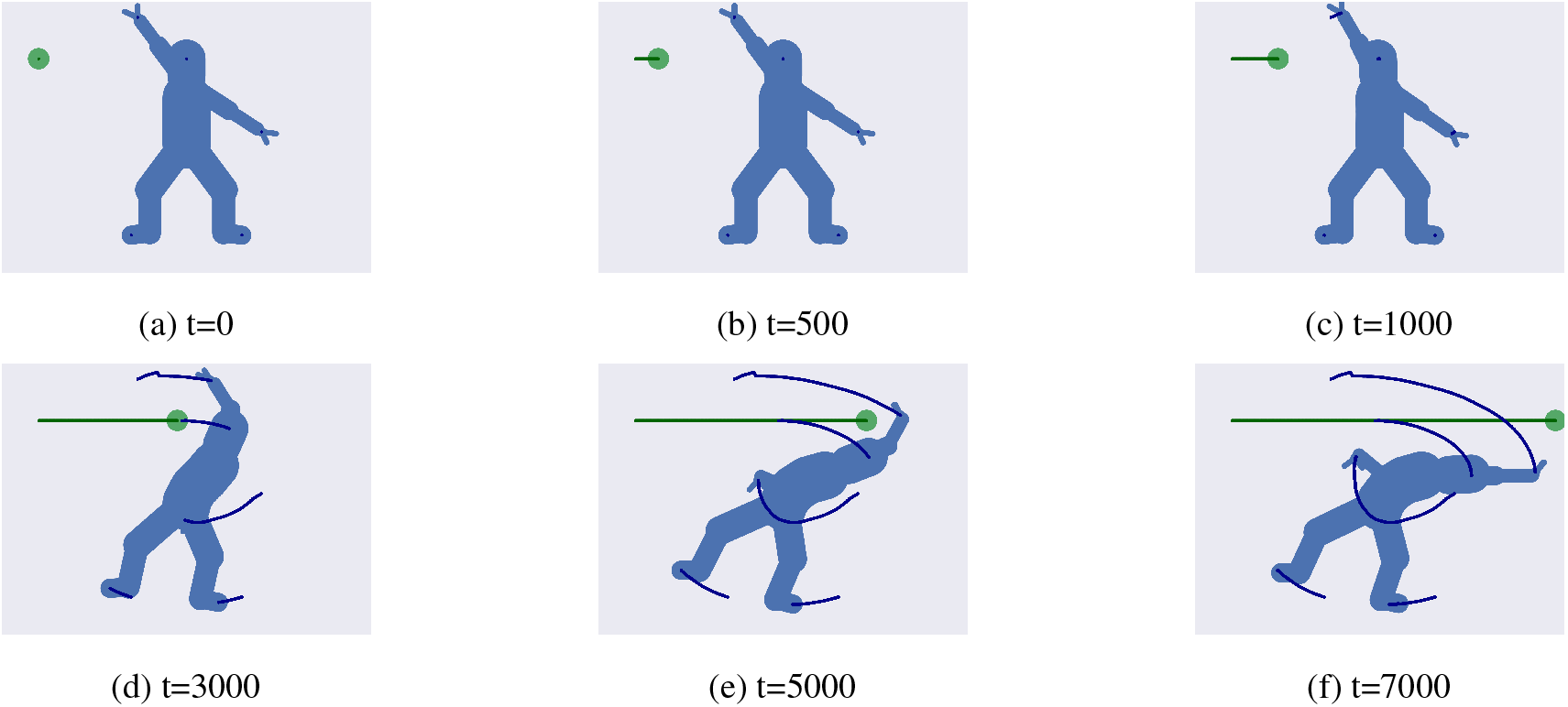
Controlling a simplified human body composed of 23-DoF. The task consists in Avoiding a dynamic obstacle with the whole body. Also see Supplementary Movie 6.

### 6.2 Videos

#### 6.2.1 Supplementary Video 1

Control of tracking (reaching a moving target, red dot) while avoiding a dynamic obstacle (green dot). Also see Supplementary Figure 8.

#### 6.2.2 Supplementary Video 2

Control of a reaching task (an extrinsic goal) while maintaining the same orientation with the last segment (an intrinsic goal, e.g., to implement a specific affordance). Also see Supplementary Figure 9.

#### 6.2.3 Supplementary Video 3

Trajectory planning in the Cartesian domain. The task consists of performing a circular trajectory. Also see Supplementary Figure 12.

#### 6.2.4 Supplementary Video 4

Controlling a complex Kinematic agent with 28 DoF. The goal consists in reaching n=4 different target locations. Also see Supplementary Figure 13.

#### 6.2.5 Supplementary Video 5

Controlling a simplified human body with 23 DoF. The task consists of reaching n=3 different target locations, one with the left knee and the other two with each of the two arms. Also see Supplementary Figure 14.

#### 6.2.6 Supplementary Video 6

Controlling a simplified human body with 23 DoF. The task consists in avoiding a dynamic obstacle with the whole body. Also see Supplementary Figure 15.

